# External mechanical cues reveal core molecular pathway behind tissue bending in plants

**DOI:** 10.1101/2020.03.05.978296

**Authors:** Anirban Baral, Emily Morris, Bibek Aryal, Kristoffer Jonsson, Stéphane Verger, Tongda Xu, Malcolm Bennett, Olivier Hamant, Rishikesh P. Bhalerao

## Abstract

Tissue folding is a central building block of plant and animal morphogenesis. In dicotyledonous plants, hypocotyl folds to form hook after seedling germination that protects their aerial stem cell niche during emergence from soil. Auxin response factors and auxin transport are classically thought to play a key role in this process. Here we show that the microtubule-severing enzyme katanin contributes to hook formation. However, by exposing hypocotyls to external mechanical cues mimicking the natural soil environment, we reveal that auxin response factors ARF7/ARF19, auxin influx carriers, and katanin are dispensable for apical hook formation, indicating that these factors primarily play the role of catalyzers of tissue bending in the absence of external mechanical cues. Instead, our results reveal the key roles of the non-canonical TMK-mediated auxin pathway, PIN efflux carriers and cellulose microfibrils as components of the core pathway behind hook formation in presence or absence of external mechanical cues.

## Introduction

Tissue folding is an essential feature of body plan development across kingdoms. In animals, tissue rearrangements can generate patterns of tension and compression, which in turn contribute to tissue invagination during gastrulation, through the actomyosin mediated response to forces (Sherrard, et al., 2010; Martin, et al., 2009; Farge, 2003). Interestingly, such intrinsic mechanical cues can be artificially mimicked by external mechanical cues in the form of local indentations; even rescuing tissue invagination defects in *Drosophila* myosin mutants (Pouille, et al., 2009). This calls for an investigation of the relative contribution of intrinsic and external mechanical cues to development. In animals the core developmental plan is laid out in the embryonic stage, and since animal embryos are usually embedded in a protective shell, such external cues are likely to be filtered out. In contrast, the postembryonic development of plants is constantly exposed to both internal and external mechanical constraints Here we investigate how plants cope with such conflicting cues to control tissue folding.

The presence of stiff cell walls and high turgor pressure in plants limit certain mechanisms of differential growth such as cell migration and cell contraction. However, because plant cells are tightly attached to one another through their contiguous walls, local differences in cell elongation is employed to create pronounced deformations in tissue and organ shape (Echevin, et al., 2019). In keeping with their sessile lifestyle, land plants have also acquired the unique ability to substantially alter their shape and form to adapt to their environment. The genetic and biochemical pathways that regulate differential growth in plants, and their modulation by exogenous cues, have been investigated in numerous studies; reviewed in (Chaiwanon, et al., 2016). In addition to biochemical signals, a critical role for mechanical signals in regulating plant development is emerging. Touch or wind has a genome-wide impact on the transcriptome, leading to wall stiffening and reduced stature (Braam, 2005) Mechanical signals can also arise internally: turgor pressure puts walls and tissues under tension (Kutschera and Nikas, 2007), and mechanical conflicts arise between cells and tissues with different growth parameters, as seen in shoot apical meristems (Uyttewaal, et al., 2012), sepals (Hervieux, et al., 2017) and leaves (Huang, et al., 2018).

Hypocotyl bending to form an (apical) hook is facilitated by differential cell elongation. Polar transport-mediated accumulation of auxin on one side of the hypocotyl restricts the elongation of epidermal cells (Abbas, et al., 2013), forming the concave side of the apical hook. Such tissue folding reflects a mechanical conflict across the structure (Landrein, et al., 2015). Interestingly, the apical hook forms during germination in the absence of external constraints in petri dishes, while under native growth conditions, hook development proceeds under mechanical constraints imposed by the soil (Abbas, et al., 2013). We are using tissue bending in the hypocotyl during seed germination as an experimental model to study how exposure to exogenous cues from mechanical constraints during soil emergence as well as endogenous cues can modulate shape change through differential growth and tissue folding.

In plants, cortical microtubules (CMTs) are key players in morphogenesis; they mediate the deposition of cellulose microfibrils and confer directionality during cell elongation. Interestingly, CMTs have been shown to align with directions of maximal tensile stress that tissue shape and/ or growth imposes (Hamant, et al., 2019; Sampathkumar, et al., 2014; Uyttewaal, et al., 2012). However, exogenously applied force can override the endogenous stress pattern and reorient CMTs (Robinson and Kuhlemeier, 2018; Uyttewaal, et al., 2012). The protein katanin plays a key role in promoting CMT self-organization in response to such cues, by severing nascent microtubule branches and microtubule crossovers as well as promoting microtubule bundling (Luptovciak, et al., 2017). In loss-of-function katanin mutants, CMT arrays are slow to self-organize in parallel bundles and thus usually appear disorganized (Bouquin, et al., 2003). CMTs in these mutants react slowly to both endogenous and exogenous mechanical cues (Hervieux, et al., 2017; Sampathkumar, et al., 2014; Uyttewaal, et al., 2012). However, attaining anisotropic CMT orientation even in the absence of katanin activity is theoretically possible through growth, cell shape and geometry-induced self-organization of CMTs (Chakrabortty, et al., 2018) and from directional stabilization of CMT arrays by prolonged exogenous mechanical cues (Hamant, et al., 2019). It seems therefore that exogenous and endogenous mechanical stress can act in synergy to control CMT organization. However, the significance of this balance of growth processes remains to be demonstrated.

Through the study of hypocotyl bending during hook formation under mechanical constraints imposed by soil, we identified a role for dynamic rearrangement of CMTs in tissue folding, in response to both internal and external mechanical cues. Importantly, our results demonstrate that external mechanical constraints can rescue tissue folding defects in the katanin mutants. The associated mechanism is dependent on cellulose microfibril organization and on the polar auxin transport machinery, and it requires an alternative auxin response pathway mediated via the TMK receptor kinase to facilitate tissue bending and apical hook formation.

## Results

### Apical hook formation depends on katanin-dependent microtubule organization

To assess the mechanisms underlying tissue bending under mechanical constraints we used apical hook development as a model. Since CMTs are key morphogenetic regulators, we first assessed their role in hypocotyl bending and hook development. We compared hook formation in WT and the *ktn1-5* mutant (Lin, et al., 2013) which lacks the microtubule-severing enzyme katanin. Whereas WT seedlings formed closed apical hooks approximately 30 Hrs after germination (Figure 1A, left), *ktn1-5* mutant seedlings could not achieve hypocotyl bending and their hooks were open (Figure 1A, right). Using time-lapse infrared imaging (Boutte, et al., 2013) we observed that *in vitro-grown* WT seedlings formed closed apical hooks by progressive bending the hypocotyl within 24-26 Hrs of germination. After 28 Hrs, hooks of all the WT seedlings were closed (hook angle of 180°) (Figure. 1B). In contrast, *ktn1-5* seedlings growing on the agar surface were unable to form this bend and 28 Hrs after germination their hook angles were much smaller (53.89 ± 25.63°, n = 20), (Figure. 1B)

**Figure. 1.**
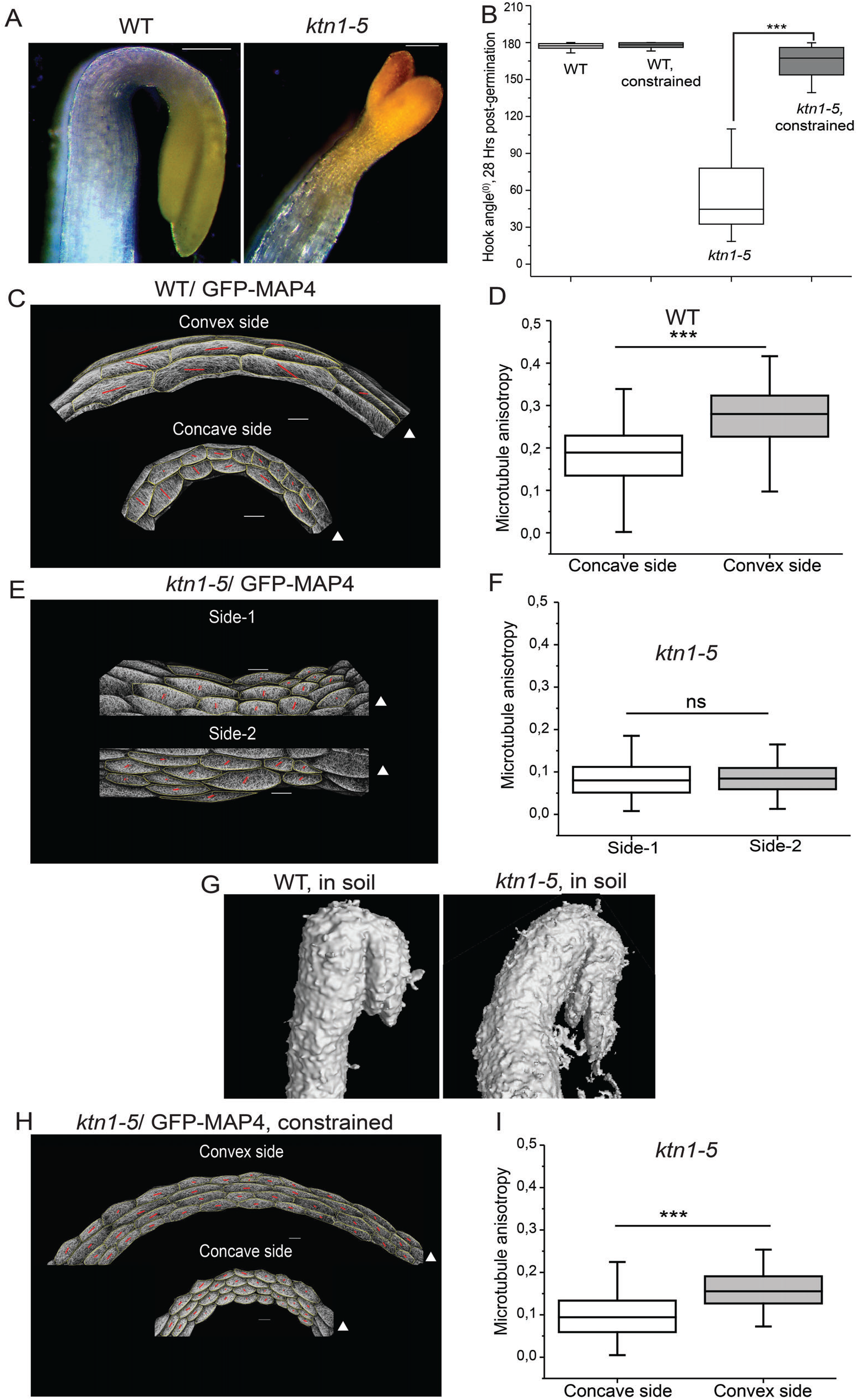
*ktr>1-5* mutant has a strong apical hook defect, which can be rectified by mechanical constrain of soil. (A) Representative image of WT and *ktn1-5* seedling grown without mechanical constrain 30 Hrs after germination. (B) Quantification of apical-hook angles in WT and *ktn1-5* seedlings growing on the surface of MS agar or growing through the gel (constrained), 28 Hrs after germination, n = 20 seedlings. (C-F) CMT organization in the hook region (0-400 μm from the shoot apical meristem) of WT, and *ktn1-5* seedlings.Representative image of CMTs visualized with GFP-MAP4 (C, E). The length of the red lines in each cell represent the average CMT anisotropy of the cell quantified by Fibriltool. Triangles denote the end of the hypocotyl with the shoot apical meristem. Quantification of CMT anisotropy values (D, F) across two sides of the hypocotyl measured by FibrilTool. Anisotropy values from at least 150 cells from 5 seedlings were quantified per group. (G) Representative image of soil embedded WT and *ktn1-5* seedling. Seeds were buried under 1 cm of sandy loamy soil and imaged by micro CT, 6 days after sowing. (H-l) CMT organization in the hook region of *ktn1-5* seedlings grown in presence of mechanical constrain (inside agar); representative image (H) and anisotropy calculation using Fibriltool (I). The data in C, D, F, and I are plotted as box plots showing the first and third quartiles, split by the median; whiskers extended to minimum and maximum values. Asterisks (***) indicate P < 0.0001 (Student’s two tailed t-test). Scale bar = 200 μm in A and 20 μm in D, E, F.

CMT organization has been associated with stem curvature in response to gravistimulation or forced bending (Ikushima and Shimmen, 2005). Here we characterized CMT arrangement in WT hooks compared to *ktn1-5.* We used the reporter microtubule-associated protein 4 fused with green fluorescent protein (GFP-MAP4) to visualize microtubules (Marc, et al., 1998) and the FibrilTool plugin (Boudaoud, et al., 2014) to quantify CMT arrangement (array anisotropy) across the hypocotyl. In WT hooks CMT arrays were significantly more anisotropic (anisotropy value 0.27 ± 0.009 SEM, n = 250 cells) on the convex side of the hook than on the concave side (anisotropy value 0.18 ± 0.001 SEM, n = 251 cells) (Figure. 1C, 1D). In contrast, CMT arrays in *ktn1-5* showed significant reduction in anisotropy and there was little difference in array anisotropy across the two sides of the hypocotyl (Figure. 1E, 1F, side1 *vs* side2; anisotropy values 0.084 ± 0.003 SEM, n = 221 cells and 0.085 ± 0.002 SEM, n = 179 cells). These results suggest that the difference in CMT organization on the two sides of the hypocotyl is associated with tissue bending during hook development and its disruption is associated with defects in hook formation.

### Apical hook defect in *katanin mutant* can be rescued by external constraints

Typically, upon germination, emerging seedlings are exposed to physical, biochemical and biological constraints by the soil. Hence we investigated the role of microtubules in hook formation in wild type and *ktn1-5* under natural conditions by germinating seeds within sterile soil and imaging hook formation using non-invasive X-ray microcomputed tomography (Orman-Ligeza, et al., 2018). Intriguingly, hypocotyl bending and formation of closed hooks were observed in *ktn1-5* seedlings growing through the soil (Figure. 1G). Thus, despite the inability to form a hook on the agar surface, during growth in soil (a more natural condition for germinating seeds), the *ktn1-5* hypocotyl was able to bend and form a hook.

To focus on the physical contribution of the soil to this response, we investigated hook formation by germinating seeds inside agar (as a proxy for soil). Using time-lapse infrared imaging (Boutte, et al., 2013), we observed that hooks of all the WT seedlings were closed (hook angle of 180^0^) both on the surface and in agar. Hook angles of constrained (agar-grown) and free (surface-grown) WT seedlings were similar, suggesting that mechanical constraints do not amplify hook bending (Figure. 1B). Intriguingly, germination in agar significantly alleviated hook defects in *ktn1-5* seedlings and after 28 Hrs post-germination their hook angles were much larger (162.06 ± 17.8°, n= 20), when compared to surface-grown *ktn1-5* seedlings, i.e. those not exposed to external mechanical constraints (Figure. 1B).

### *katanin* rescue by external constraints relies on microtubule dynamics

The developmental defects seen in the *ktn1-5* mutant are attributed to the CMT disorganization associated with loss of the microtubule-severing and bundling activities of katanin (Luptovciak, et al., 2017). In the most parsimonious scenario, reestablishment of apical hook formation by mechanical cues in *ktn1-5* could suggest that microtubules are not essential for bending in response to mechanical cues during growth in soil. To check this hypothesis, we used pharmacological and genetic approaches. Treatment with low concentrations of the microtubule-depolymerizing drug oryzalin (200 nM) or the microtubule-stabilizing drug paclitaxel (50 nM), which did not affect WT seedlings (Figure. S1A, S1C), completely abolished hook formation in *ktn1-5* seedlings grown in agar (Figure. S1B, S1D). Additionally, genetically perturbing CMT rearrangement by introducing the *tua5*^D251N^ mutation (which slows down microtubule dynamics) (Ishida, et al., 2007) in *ktn1-5* also abolished hook formation in response to mechanical constraints (Figure. S1F). Thus, even in the absence of katanin, microtubule dynamics is required for apical hook formation.

When mechanical stress levels are increased in the shoot apical meristem (e.g. upon wall weakening with the cellulose synthase inhibitor isoxaben), CMTs can exhibit WT-like behavior in a *katanin* mutant, where they can reorient around a single cell ablation as in WT (Uyttewaal, et al., 2012). We reasoned that soil, or agar gel, may provide a mechanical cue strong enough to compensate for the slower CMT dynamics in the *katanin* mutant. To test this hypothesis, we investigated whether constrained growth in agar could facilitate a difference in CMT organization in the *ktn1-5* mutant. In contrast to the lack of significant difference in CMT array anisotropy in surface-grown *ktn1-5* seedlings, growth under mechanical constraints resulted in a significant difference in CMT array anisotropy between the two sides of *ktn1-5* mutant hypocotyls (Figure. 1H, 1I; anisotropy value of 0.15 ± 0.003 SEM, n = 315 cells on convex side *vs* 0.09 ± 0.002 SEM, n = 360 cells on concave side). Note that absolute anisotropy values were lower in *ktn1-5* than in the WT, suggesting that the relative difference in CMT behavior between the two sides of the hypocotyl may be sufficient for tissue bending. Taken together our results indicate that external mechanical cues can catalyze the formation of differential CMT organization on both sides of the hypocotyl to induce tissue bending.

### Cellulose microfibril deposition is critical for mechanically-induced hook formation

From the close association of CMTs and cellulose synthase complexes (Paredez, et al., 2006) it can be surmised that CMTs would guide the deposition of cellulose microfibrils to promote hook formation. To formally check this, we first examined hook formation in two mutants with altered cellulose content and cellulose organization: the cellulose synthase A1 (CesA1) mutant *any1*, with reduced crystalline cellulose content (Fujita, et al., 2013), and the cellulose synthase interacting 1 (CSI1) protein deficient mutant *pom2-8* (Bringmann, et al., 2012), featuring altered cellulose fiber orientation arising from the partial decoupling of CesA complex from CMTs. As expected, both *any1* and *pom2-8* mutants had defects in hook formation (Figure 2A, 2B). However, unlike *ktn1-5,* hook formation in *any1* and *pom2-8* mutants was not appreciably suppressed by growth inside agar (Figure. 2A, 2B). These results further suggest that the rescue of the *ktn1-5* hook defect by growth in agar requires CMT-guided cellulose deposition. To test this, we introduced *any1* and *pom2-8* mutations into the *ktn1-5* background. Hook formation could not be restored in agar-grown *ktn1-5 any1* or *ktn1-5 pom2-8* double mutant seedlings (Figure. 2C, 2D). We also found that hook formation in agar-grown *ktn1-5* seedlings was hypersensitive (Figure. S2) to 100 nM 2,6-dichlorobenzonitrile (DCB), a cellulose biosynthesis inhibitor (DeBolt, et al., 2007). Hence, both genetic and pharmacological approaches demonstrate the crucial role of cellulose organization in mediating the bending required for hook formation in response to mechanical cues.

**Figure. 2.**
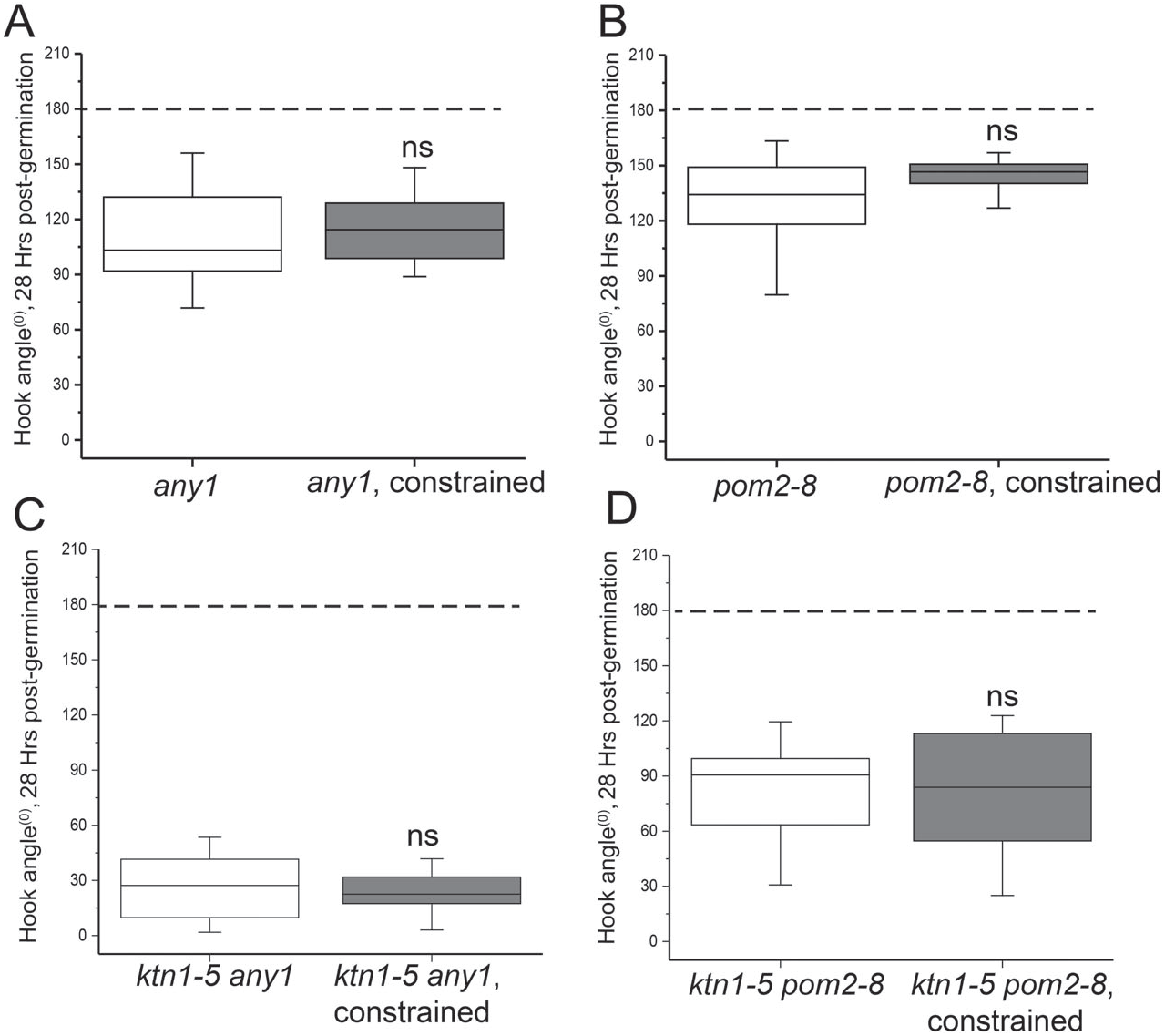
Hook formation defects in cellulose organization mutants is not rectified by mechanical constrain. Quantification of apical-hook angles in *any1* (A) and *pom2-8* seedlings (B) growing on the surface of MS agar or growing through the gel (constrained), 28 Hrs after germination, n = 20 seedlings. (C-D) Quantification of apical hook angles in *ktn1-5 any1* (C) and *ktn1-5 pom2-8* (D) double mutant seedlings. 28 Hrs after germination, n = 16 seedlings. The data in is plotted as box plots showing the first and third quartiles, split by the median; whiskers extended to minimum and maximum values. The dashed line denotes hook angle of 180°. ns = non-significant at P ≤ 0.05 (Student’s two tailed t-test).

### Differential growth mediated by mechanical cues is dependent on auxin response asymmetry

Hypocotyl bending during apical hook formation results primarily from differences in epidermal cell elongation between the sides (Raz and Koornneef, 2001). Consistent with this, and because there is limited cell division at this stage (Zadnikova, et al., 2016), epidermal cell length was significantly different between the concave (34.26 ± 0.11 μm SEM, n = 250 cells) and convex (61.39 ± 2.86 μm SEM, n = 251 cells) sides of the WT hook (Figure 3A). In sharp contrast, epidermal cell lengths in *ktn1-5* were similar on both sides of the hypocotyl (Figure. 3B, side 1 vs side 2); cell length 53.85 ± 1.592 μm SEM, n = 221 cells and 61.90 ± 2.02 μm SEM, n = 179 cells. As expected, when grown inside agar, cell length difference across the hypocotyl was re-established in *ktn1-5* seedlings (Figure. 3C); cell length of 33.02 ± 0.64 μm SEM, n = 360 cells on the concave side *vs* 63.24 ± 1.2 μm SEM, n = 351cells on the convex side. The cells on the concave (inner) side of mechanically constrained *ktn1-5* seedlings were significantly smaller than cells on either side of surface-grown *ktn1-5* seedlings (Figure. 3D). These results suggest that growth asymmetry mediated by repression of growth on one side can be reestablished in *ktn1-5* seedlings grown under mechanical constraint.

**Figure. 3.**
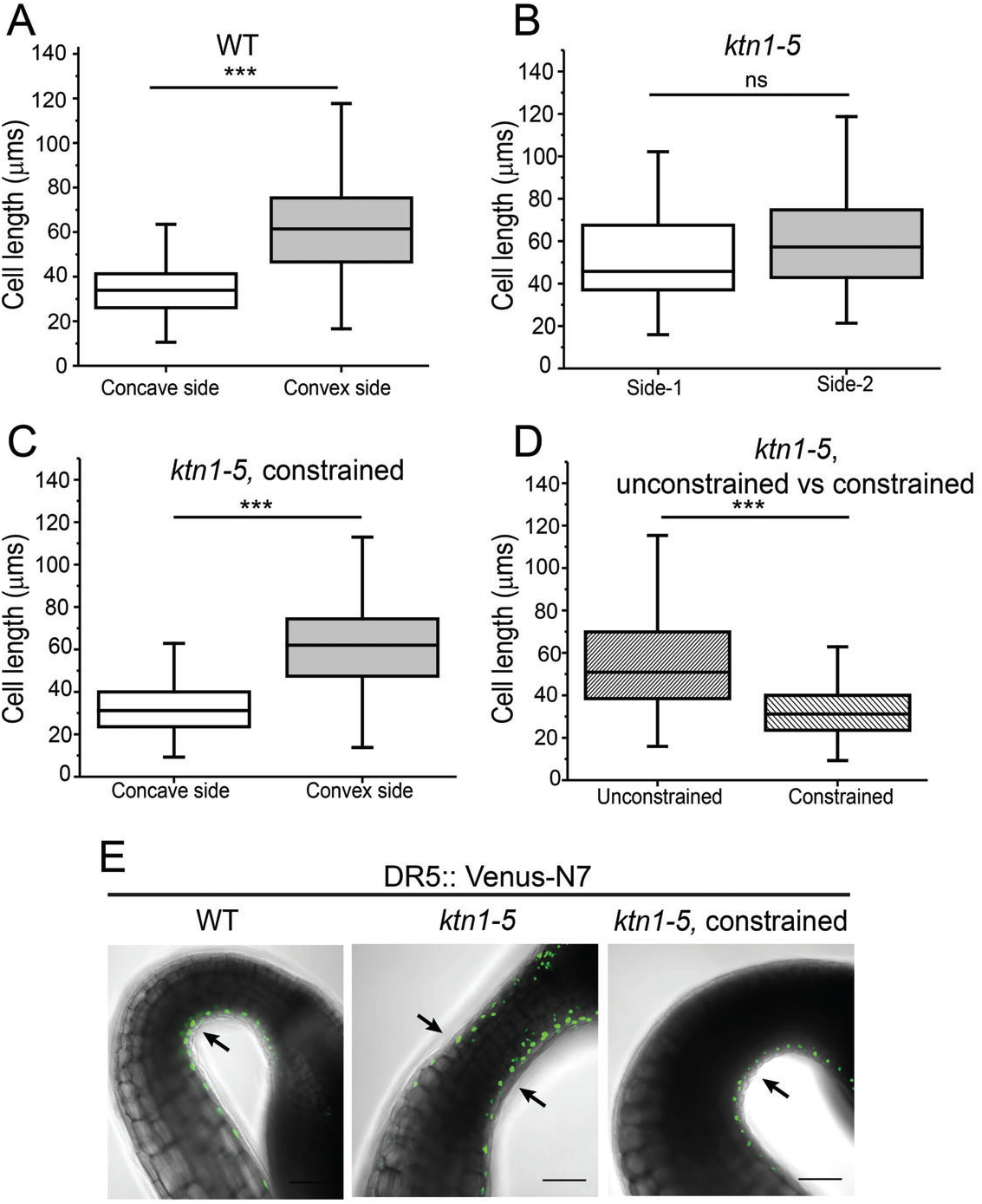
Mechanical constrain reestablishes auxin response maxima and cell length repression in *ktn1-5* seedlings. (A-C) Quantification of cell length of WT (A), *ktn1-5* (B) and mechanically constrained *ktn1-5* seedlings (C) in the hook region (0-400 μm from the shoot apical meristem). Cell length is measured from at least 150 cells from 5 seedlings per group, where cell outlines were marked with GFP-MAP4. (D) Comparison of average cell length in the hook region of unconstrained *ktn1-5* seedlings with concave side cells constrained *ktn1-5* seedlings shows cell growth repression reestablishment by mechanical constrain. Box plots show the first and third quartiles, split by the median; whiskers extended to minimum and maximum values. Asterisks (***) indicate P < 0.0001, ns = non-significant at P ≤ 0.05 (Student’s two tailed t-test). (E) Representative image of DR5:: Venus-N7 signal in WT and *ktn1-5* seedlings, 28-30 Hrs post germination. In WT, DR5:: Venus-N7 signal is detected only in epidermal cells on the concave side of apical hook (arrow). In *ktn1-5,* DR5::Venus-N7 signal is seen on both sides of the hypocotyl (arrows) suggesting a defect in auxin asymmetry establishment. DR5::Venus-N7 signal asymmetry is reestablished in *ktn1-5* by mechanical constrain by growing inside agar. Scale bar = 100 μm.

The repression and slower rate of cell elongation can be attributed to auxin response maxima forming on the concave side (Zadnikova, et al., 2010). Mechanical constraints in agar-grown hypocotyls could simply deform the hypocotyl resulting in bending through a direct impact on CMTs. Alternatively, mechanical cues may influence the auxin response, modifying differential growth, and indirectly affecting CMTs. To address this, we first investigated whether impaired hook formation in *ktn1-5* seedlings is due to attenuation of the auxin response. In WT hooks, auxin response maxima, visualized using the synthetic auxin reporter DR5:: Venus-N7 (Heisler, et al., 2005), were seen only on the concave side (Figure. 3E, left): whereas the *ktn1-5* seedlings failed to establish this asymmetry; DR5:: Venus-N7 signals were seen on both sides of the hypocotyl (Figure. 3E, middle). We then investigated whether constrained growth in agar gel could reestablish DR5:: Venus-N7 signal asymmetry in a katanin-independent manner. In *ktn1-5* seedlings grown inside agar, the DR5:: Venus-N7 signal was observed only on the concave side (Figure. 3E, right). Lastly, microtubule perturbation with 200 nM oryzalin abolished establishment of DR5:: Venus-N7 signal asymmetry in *ktn1-5* seedlings grown inside agar (Figure. S3B). These data suggest that mechanical constraints in agar-grown hypocotyls induce hook formation by establishing auxin response asymmetry, and the resulting mechanical conflict would be strong enough to override reduced CMT dynamics in the *katanin* mutant.

### Mechanical constraints can modulate polar auxin transport during apical hook formation

To decipher the link between external mechanical constraints and differential auxin response in the apical hook, we next analyzed the contribution of auxin transporters. Auxin influx carriers of the AUX/ LAX family are thought to play an important role in auxin response asymmetry in the apical hook (Vandenbussche, et al., 2010). Consistently, hook formation was severely affected in the *aux1 lax1 lax2 lax3* quadruple mutant when grown on the surface, mimicking the *ktn1-5* mutant phenotype (Figure. S4A). As observed for *ktn1-5,* growth inside agar could restore hook formation in this *aux/lax* quadruple mutant (Figure. S4A), as observed for *ktn1-5.* Thus hook formation does not require AUX/LAX auxin influx carriers.

The efflux carriers of the PIN-FORMED (PIN) family are also thought to be required for hook formation (Zadnikova, et al., 2010). Similar to AUX/ LAX carriers, hook formation was also completely abolished in the *pin1347* quadruple mutant when grown on the surface (Figure. S4B). However, in contrast with *aux/ lax* or *ktn1-5* mutants, hook formation did not occur in *pin1347* (Figure. S4B) even when grown inside agar. Thus external mechanical cues can override both katanin-dependent CMT dynamics and AUX/ LAX-dependent auxin influx, but they require PIN auxin transporters for hook formation. Tissue- and cell-type-specific polar localization of PIN transporters is critical for the directional flow of auxin (Grunewald and Friml, 2010). For the PINs implicated in hook development (PIN1,3, 4 and 7), PIN4-GFP signal could be detected only in the epidermal cells. The localization of PIN4-GFP was strongly polar with preferential localization to transverse membranes compared to the lateral sides (Figure 4A, left; Figure. S5A, S5B). PIN3-GFP signal could be detected in epidermal and cortical cells, with significantly stronger signal on transverse membranes compared to lateral sides in the epidermis (Figure 4B, left; Figure. S5A, S5C) and exclusively transverse localization in cortical cells (Figure 4C, left). In surface-grown *ktn1-5* mutant seedlings, the polarities of PIN3-GFP and PIN4-GFP were strongly perturbed. Localization of both PIN3 and PIN4 GFP was significantly enhanced at the lateral membranes of epidermal cells in *ktn1-5* compared to WT (Figure. 4A, 4B, middle; Figure. S5B, S5C). Moreover, PIN3-GFP was mislocalized to lateral membranes in cortical cells of *ktn1-5* (Figure 4C, middle). Intriguingly, PIN1-GFP displayed a unique spatially-restricted localization pattern in the WT hook (Zadnikova, et al., 2010). PIN1-GFP signal could be detected on the plasma membrane only in cells on the concave side (Figure 4E, left). In the *ktn1-5* hook region, however, PIN1-GFP was completely absent from plasma membrane but it could be detected in punctate intracellular structures (Figure 4E, middle). PIN7-GFP signal could be detected only in epidermal cells, which showed nonpolar localization (Figure S5). In contrast to PIN 1,3 and 4, neither the localization nor the intensity of PIN7 underwent any change in *ktn1-5* (Figure S6D, S6E). We then examined whether mechanical cues can mediate the localization of PIN transporters required to induce hook formation. When grown inside agar, the polarity of PIN4-GFP in epidermis (Figure 4A right; Figure. S6B) and PIN3-GFP in both epidermis (Figure 4B, right; Figure. S6C) and cortex (Figure. 4C, right) was partially but significantly restored in *ktn1-5.* Furthermore, PIN1-GFP signal was now primarily on the plasma membrane of cells on the concave side in *ktn1-5,* resembling the localization pattern seen in WT hooks (Figure. 4D, right).

**Figure. 4.**
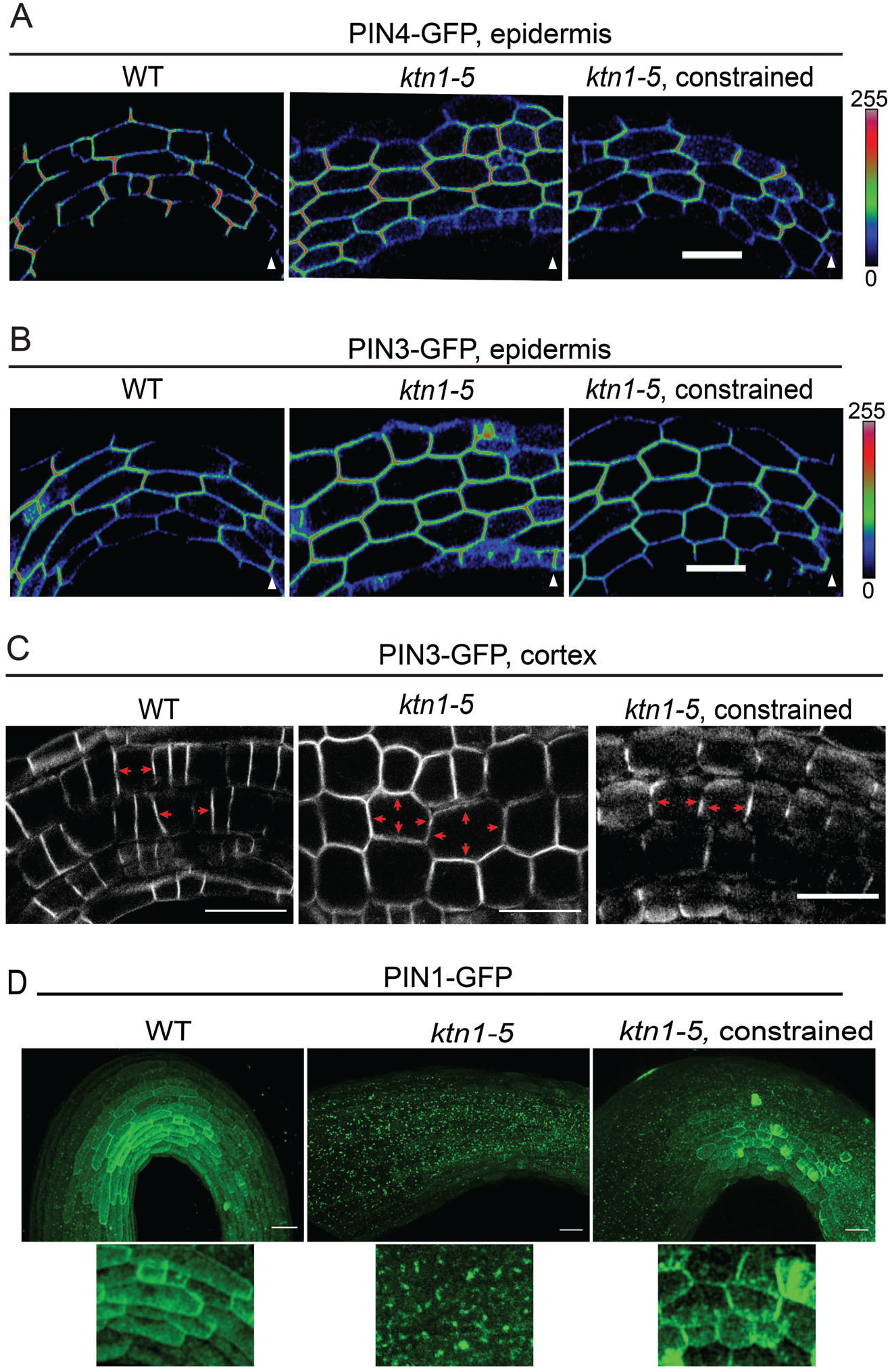
Altered localization of PIN auxin transporters in *ktn1-5* could be rectified by mechanical constrain. Representative images of PIN4-GFP in epidermis (A), PIN3-GFP in epidermis (B) and PIN3-GFP in cortex (C) in WT, *ktn1-5* and mechanically constrained *ktn1-5* seedlings. (D) PIN1-GFP in WT hook (left) shows plasma membrane localization only on concave side of the hook. In *ktn1-5* seedlings PIN1-GFP is absent from the plasma membrane (middle). In mechanically constrained *ktn1-5* seedlings, PIN1-GFP is seen on plasma membrane on cells on concave side (right). Magnified images of concave side cells are shown underneath. Scale bar = 50 μm. Color bars in (A) and (B) represent 8-bit intensity range of 0-255, triangles denote the end of the hypocotyl with the shoot apical meristem. See also **Figure. S5**.

Thus, the genetic analysis of *pin* mutants and localization analysis of PINs strongly supports our scenario: mechanical constraints associated with growth in agar would mediate the recruitment (PIN1) and polarity (PIN3, PIN4) of auxin efflux carrier, thereby generating a differential auxin response and differential cell elongation; the resulting mechanical conflicts would be strong enough for CMTs to align with the directions of mechanical stress, and maintain tissue bending, even compensating the absence of the catalyzer of CMT self-organization, katanin.

### Cellulose microfibril deposition is critical for differential auxin response in the hook

As mentioned above, cellulose acts downstream of CMTs to control directional cell growth, through the CMT-CesA nexus (Paredez, et al., 2006). Interestingly, a role for CesA activity in the regulation of PIN1 localization has also been reported (Feraru, et al., 2011). This would put cellulose upstream of the CMT response to mechanical cues, via the formation of differential auxin patterns and resulting mechanical conflicts. This prompted us to investigate whether external cues require cellulose microfibrils for the establishment of PIN polarities and auxin response. First, we investigated whether the cellulose mutants fail to establish the auxin response maxima critical for hook formation. In contrast to WT plants, with DR5:: Venus-N7 signal only on the concave side, Figure. 5A), the DR5:: Venus-N7 pattern in seedlings of both the cellulose mutants, *any1* and *pom2-8*, was altered and diffuse (Figure. 5B, 5C) suggesting a defect in establishing an auxin response maximum when cellulose deposition is affected. Analysis of the localization of PIN efflux carriers showed epidermal polarity of PIN3-GFP and PIN4-GFP was significantly reduced in *any1* and *pom2-8* compared to WT (Figure. 5D, 5E, S6A, S6B). PIN3-GFP was also found on lateral membranes of cortical cells in *any1* and *pom2-8* hooks (Figure. 5F), similar to *ktn1-5* seedlings. Looking at PIN1-GFP, we found that, similar to *ktn1-5,* GFP signal was absent from plasma membrane but present in intracellular punctate structures (Figure. 5G). These results are consistent with a scenario in which cellulose is required for the polarity of PIN auxin efflux carrier and the resulting auxin response patterns; in that scenario, cellulose would indirectly control the CMT response to external mechanical cues, via its impact on auxin distribution.

**Figure. 5.**
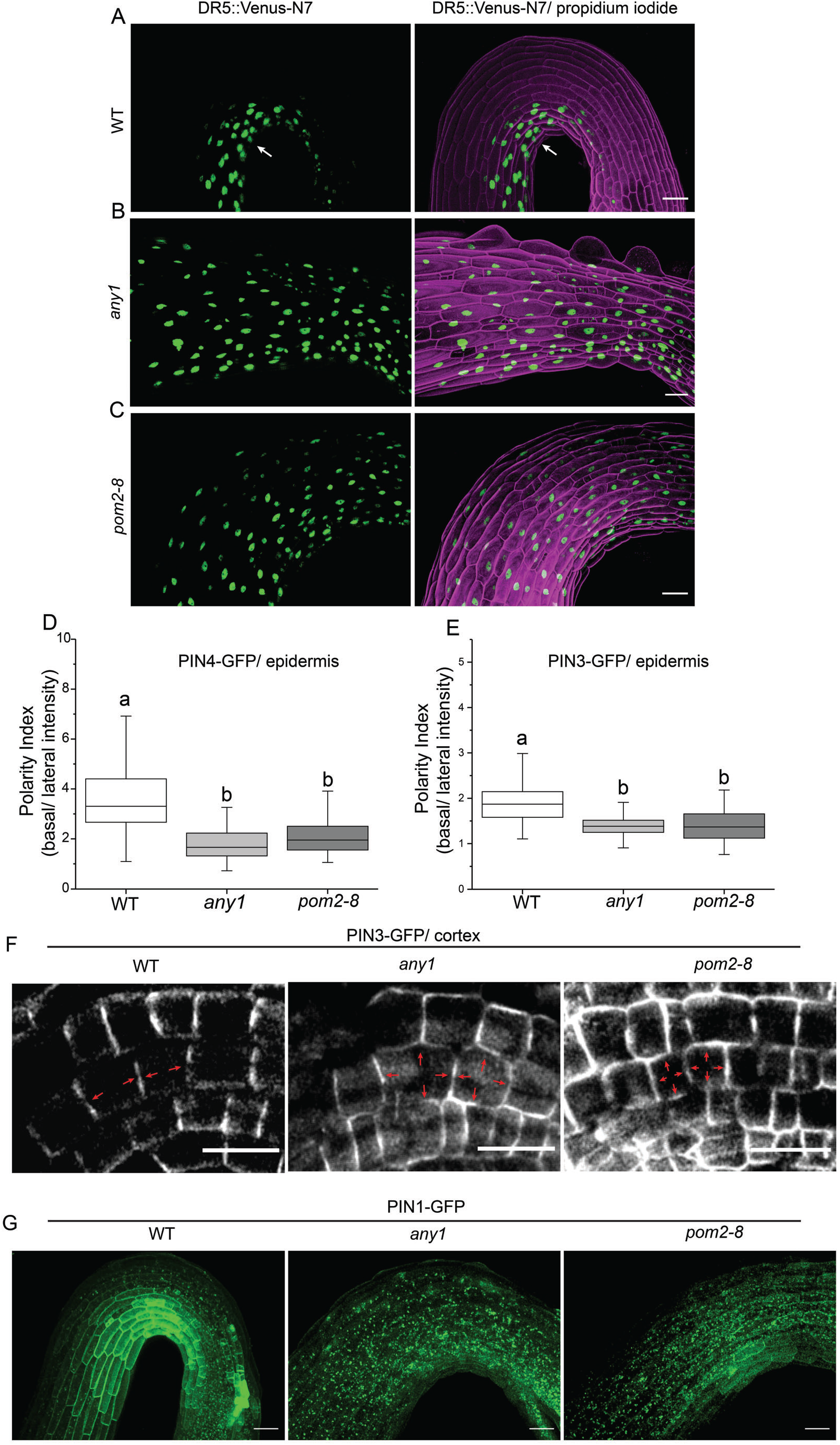
Altered auxin response asymmetry and PIN localization in cellulose mutants. (A-C) Representative image showing Z-projection of DR5:: Venus-N7 signal in WT (A), *any1* (B) and *pom2-8* (C) seedlings. The cells are counterstained with propidium iodide. The concave side specific localization of DR5::Venus-N7 signal seen in WT (arrow) is altered in *any1* and *pom2-8* Scale bar = 50 μm. (D-E) Quantification of polarity index (basal/ lateral fluorescence intensity) of PIN4-GFP (D) and PIN3-GFP (E) in epidermal cells of hook region of WT, *any1* and *pom2-8* seedlings. Polarity index was measured from at least 100 cells from 10 seedlings/ group. Box plots show the first and third quartiles, split by the median; whiskers extended to minimum and maximum values. Groups with significant difference by one-way ANOVA with post hoc Tukey’s test (P < 0.05) are indicated by different letters. (F) Representative images showing localization of PIN3-GFP in hook region cortical cells of WT, *any1* and *pom2-8* seedlings. (G) Localization of PIN1-GFP in WT, *any1* and *pom2-8* seedlings. Scale bar = 50μm. See also **Figure. S6.**

### Signaling mediated by TMK1 and 4 is necessary for mechanosensitive hook formation

Two major pathways have been reported to mediate the auxin response in the hook. The transcription factors ARF7/ ARF19 mediate the canonical auxin response pathway, and as a result the *arf7 arf19* double mutant displays a strong hook defect (Zadnikova, et al., 2010). However, we found that the marked hook defects of the *arf7 arf19* double mutant could be rescued by constraining growth in agar gel (Figure. S7A). Furthermore, the *ktn1-5 arf7 arf19* triple mutant displayed a similar response to mechanical cues as the *ktn1-5* single mutant with respect to hook formation (Figure. S7B). This suggests that hook formation in response to mechanical cues does not require ARF7 and ARF19 activity.

A recent study suggests that the plasma membrane localized receptor TMK1 is a component of a non-canonical auxin signaling pathway operating in the apical hook (Cao, et al., 2019). Perception of higher levels of auxin causes cleavage and nuclear translocation of a C-terminal fragment of TMK1 specifically on the concave side of the hook, subsequently repressing growth of these cells through stabilization of specific nuclear AUX/ IAA transcription factors. Since neither ARF7 and ARF19 was required for hook formation in response to external mechanical cues, we investigated whether this pathway acts via TMKs.

The *tmk1* null mutant displays a defect in hook maintenance but not in hook formation. Nor do the loss of three remaining members of the TMK family cause any appreciable hook phenotypes (Cao, et al., 2019). However, the functional redundancy between TMK1 and TMK4 (Dai, et al., 2013) prompted us to look into hook formation in the *tmk1tmk4* double mutant. This mutant showed a clear hook formation defect, as well as defects in auxin response asymmetry as visualized with DR5:: Venus-N7. Strikingly, this response could not be rescued by constraining growth in agar gel (Figure. 6A-C). These results strongly suggest that hook formation in response to growth under mechanical constraints depends on the TMK pathway. Taken together, our observations suggest that the components of the non-canonical auxin response pathway, TMK1 and TMK4, act redundantly and are required to mediate bending in response to mechanical cues during hook formation.

**Figure. 6.**
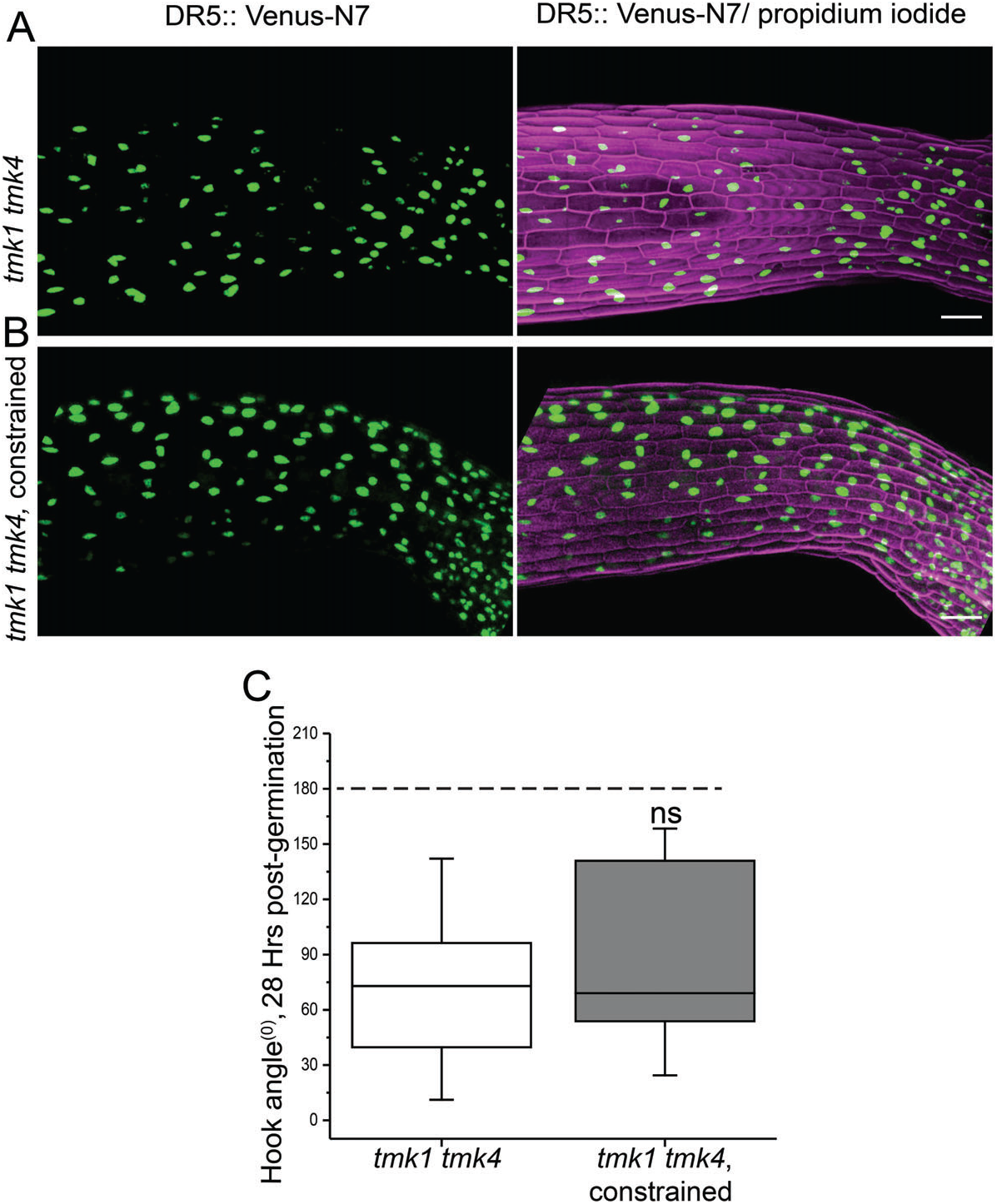
TMK1 and TMK4 is required for hook formation under mechanical constrain. Representative image showing Z-projection of DR5::Venus-N7 signal in *tmk1 tmk4* seedling in absence (A) and in presence of mechanical constrain (B). The cells are counterstained with propidium iodide, Scale bar = 50 μm. (C) Quantification of apical-hook angles in *tmk1 tmk4* growing on the surface of MS agar or growing through the gel (constrained), 28 Hrs after germination, n = 16 seedlings. Box plots show the first and third quartiles, split by the median; whiskers extended to minimum and maximum values. The dashed line denotes hook angle of 180°. ns = non-significant at P ≤ 0.05 (Student’s two tailed t-test).

## Discussion

A unique feature of morphogenesis in plants is the postembryonic adaptability to their environment. Formation of the hook during early seedling establishment is an example of such an adaptive response, protecting the fragile shoot apical meristem during soil emergence. In this study we show that asymmetry in cell elongation, the auxin response and CMT behavior drive differential growth during hook formation. Although these responses are absent in the *katanin* mutant, we demonstrate that katanin only catalyzes this response: external mechanical cues can generate a hook independent of katanin. Recently, an important role of mechanical cues originating from gravitropic bending of the root to the generation of auxin asymmetry has been proposed during hook formation (Zhu, et al., 2019). Our results provide support for this suggestion and reveal a scenario involving auxin efflux carriers, and not auxin influx carriers, and in which CMTs react to external mechanical cues and a TMK-dependent auxin response pathway to promote hook formation.

Although apical hook formation has been studied extensively as a model for differential cell elongation and underlying hormonal pathways and transcriptional networks (Mazzella, et al., 2014), most of these studies were performed by growing seeds on the surface of agar. Under these laboratory conditions, the hypocotyl is not subject to external cues akin to mechanical constraints imposed by soil. As a result, the role of exogenous mechanical cues in hypocotyl bending to form a hook has not been explored in detail. By introducing external mechanical cues in the form of agar-grown hypocotyls, we reveal that three key factors, auxin response factors, auxin influx carriers and microtubule severing enzyme katanin are dispensable for apical hook formation. By challenging the intrinsic genetic control of apical hook formation with external mechanical cues, we could extract the key players of tissue bending in the context of apical hook formation.

How plant cells may differentiate between external (e.g. wind, touch) and internal mechanical cues is subject to debate. Plant cells may in fact use the same pathways to respond to mechanical forces, independent of the cause of mechanical stress (Hamant and Haswell, 2017). In particular, artificial mechanical perturbations are classically performed to check whether plant cells can orient their CMTs with the direction of maximal tensile stress, mimicking the CMT response to growth- and shape-derived stress in control tissues (Robinson and Kuhlemeier, 2018; Sampathkumar, et al., 2014; Hamant, et al., 2008). Yet, the associated mechanotransduction pathways have remained unknown so far. Among the candidate mechanisms to exclude, CMTs seem to align with local mechanical stress reorientation in the shoot apical meristem even when calcium signals are blocked (Li, et al., 2019) or when auxin gradients are largely impaired (Heisler, et al., 2010) Therefore it has been proposed that CMTs may be their own mechanosensors (Hamant, et al., 2019) echoing what has been proposed for actin in animals (Yu, et al., 2017; Risca, et al., 2012) and building on evidence from stretched microtubules *in vitro* (Franck, et al., 2007). Yet, the integration of such behaviors in development requires a coordinator, a role that could be played by auxin and mediated by a TMK dependent, non-canonical auxin response pathway.

Several mechanisms mediate the formation of ordered CMT arrays, including polymerization/ depolymerization, bundling and severing (Dixit and Cyr, 2004). Microtubule severing by katanin has been shown to promote CMT anisotropy in several studies; reviewed in (Luptovciak, et al., 2017) and indeed our data also show that in the absence of katanin, CMT arrays are more isotropic. However, we found that mechanical cues can promote CMT anisotropy in the absence of katanin. At first glance, our observations are in conflict with the role of katanin in mediating the response of CMTs to external mechanical cues (Sampathkumar, et al., 2014). However, our data are consistent with slower but pronounced reorganization of CMTs in meristems of katanin mutants subjected to prolonged mechanical cues (Louveaux, et al., 2016; Uyttewaal, et al., 2012). Katanin-independent CMT reorganization can happen by motor protein mediated reorientation of CMTs (Inoue, et al., 2016). Also, altered polymerization/ depolymerization of MTs by mechanical stress can rearrange CMTs in absence of katanin activity (Trushko, et al., 2013). Indeed, hypersensitivity of mechanical constrain mediated hook formation in *ktn1-5* seedlings towards MT-depolymerizing and polymerizing drugs (oryzalin and paclitaxel) point towards an important role of microtubule polymerization/ depolymerization in katanin-independent CMT reorganization.

CMT disorganization associated with loss of katanin function alters the organization of overlaying cellulose microfibrils (Burk and Ye, 2002); this, together with the alteration of PIN1 polarity associated with cellulose defects (Feraru, et al., 2011), raises the possibility that the PIN localization defects seen in the *ktn1-5* mutant stems from cellulose disorganization. That PIN mislocalization is similar in *any1* and *pom2-8* mutants supports the importance of cellulose organization in guiding the PIN polarity necessary for hook formation. Furthermore, it has been reported that PIN1 localization at the plasma membrane is sensitive to mechanical stress (Nakayama, et al., 2012). It is conceivable that CMTs and cellulose regulate PIN localization and auxin asymmetry formation indirectly, by mediating mechanical stress associated with tissue folding. Nonetheless, our study reveals that CMTs and cellulose, by acting as mediators of mechanical stress, can guide key developmental processes such as tissue folding in plants by modulating PIN localization and auxin asymmetry. The localization of TMKs at the cell periphery, their role in auxin perception (Dai, et al., 2013) together with their role in apical hook formation, opens the possibility that these receptors may act as an integrator of both biochemical (auxin) and mechanical cues.

To conclude, the analysis of apical hook formation in the presence of external mechanical cues are reminiscent of observations in *Drosophila.* During gastrulation, tissue deformation through external mechanical cues can compensate defective myosin in *Drosophila* (Pouille, et al., 2009). Here we find that stiff agar can trigger apical hook formation, even when CMT dynamics is reduced in the katanin mutant. Beyond the parallel between plants and animals, such approaches further show how challenging development with external mechanical cues can be a productive way to dissect the molecular pathways behind morphogenesis, and identify the core players

## Acknowledgements

We thank Eva Benkova, Geoffrey O. Wasteneys, Ranjan Swarup and Enrico Scarpella for sharing published materials. Our sincere thanks to Anne-Lise Routier-Kierzkowska and Daniel Kierzkowski for reading through the manuscript and providing valuable inputs. A.B thanks the Wenner-Gren foundation for a travel fellowship. This work was funded by grants from the Knut and Allice Wallenberg Foundation awarded to R.P.B

## Author Contributions

A.B, R.P.B, O.H and M.B conceived the project and designed experiments. E.M performed the MicroCT experiments. B.A contributed the *auxlax* quadruple mutant data. A.B performed all other experiments and analysed the data. K.J contributed to initial screening of the *katanin* mutant phenotypes and provided valuable discussions. T.X provided critical intellectual inputs. S.V provided the Surfcut analysis tool and provided critical inputs. A.B, R.P.B, O.H and M.B wrote the paper. R.P.B guided the research.

## Declaration of Interests

The authors declare no competing interests.

## Star methods

### Plant material

*Arabidopsis thaliana* ecotype Col-0 was used as wild type (WT) control in all experiments. The mutant lines *ktn1-5* (Lin, et al., 2013), *any1* (Fujita, et al., 2013), *pom2-8* (Bringmann, et al., 2012), *tua5*^D251N^ (Ishida, et al., 2007), *pin1 3 4 7* (Belteton, et al., 2018), *aux1 lax1 lax2 lax3* (Vandenbussche, et al., 2010) *arf7arf19* (Okushima, et al., 2005) *tmk1tmk4* (Dai, et al., 2013) and fluorescent reporter lines 35S:: GFP-MAP4 (Marc, et al., 1998), PIN1:: PIN1-GFP, PIN3:: PIN3-GFP (Zadnikova, et al., 2010), PIN4:: PIN4-GFP, PIN7:: PIN7-GFP (Belteton, et al., 2018), DR5:: Venus-N7 (Heisler, et al., 2005) lines have been described previously.

### Time lapse analysis of apical hook development

For time lapse imaging of apical hook development, seegs sown on surface of agar plates or embedded in agar (1/2 Murashige and Skoog medium, 1% plant agar, Duchefa Biochemie), were grown on vertical plates in the dark at 21°C illuminated with far infrared-light (850 nm). Seedlings were photographed every hour using a Canon D50 camera without the infrared filter. Hook angles were measured as previously described (Boutte, et al., 2013) using Image J/Fiji.

### CMT imaging and quantification

Dark-grown WT or ktn1-5 mutant plants expressing GFP-MAP4 were imaged with a Zeiss LSM 880 confocal microscope using airyscan scanning. Post-acquisition, the images were processed using the airyscan processing module incorporated in the Zen 2.3 software. The cortical signal (0-10 μm depth) was extracted from the stacks using the Surfcut pipeline incorporated in the Image J/Fiji program (Erguvan, et al., 2019). CMT anisotropy was measured using the FibrilTool plugin (Boudaoud, et al., 2014) in Image J/Fiji.

### Confocal laser-scanning microscopy and quantification of fluorescence intensity

Samples were imaged using a Ziess LSM 880 confocal laser scanning microscope. GFP, and YFP, were excited by 488 and 514 lasers respectively and respective fluorescence emissions were detected at 492-540 and 518-560 nm windows. Images for plasma membrane fluorescence intensity quantification of PIN1-GFP, PIN3-GFP, PIN4-GFP and PIN7-GFP in hook epidermal cells of were captured under identical acquisition parameters (laser power, photomultiplier gain, offset, zoom, and resolution) between WT and mutants. Polarity index was calculated as the ratio of basal/ lateral intensities. Statistical analysis methods are described in figure legends.

### X-Ray microcomputed tomography

WT and *ktn1-5* seeds were buried under 1 cm of sterile, sandy loamy soil. Seedlings were scanned after 6 days post-sowing by X-Ray microcomputed tomography as described previously (Orman-Ligeza, et al., 2018) at 6.5 μm resolution.

---

**Figure. S1.**
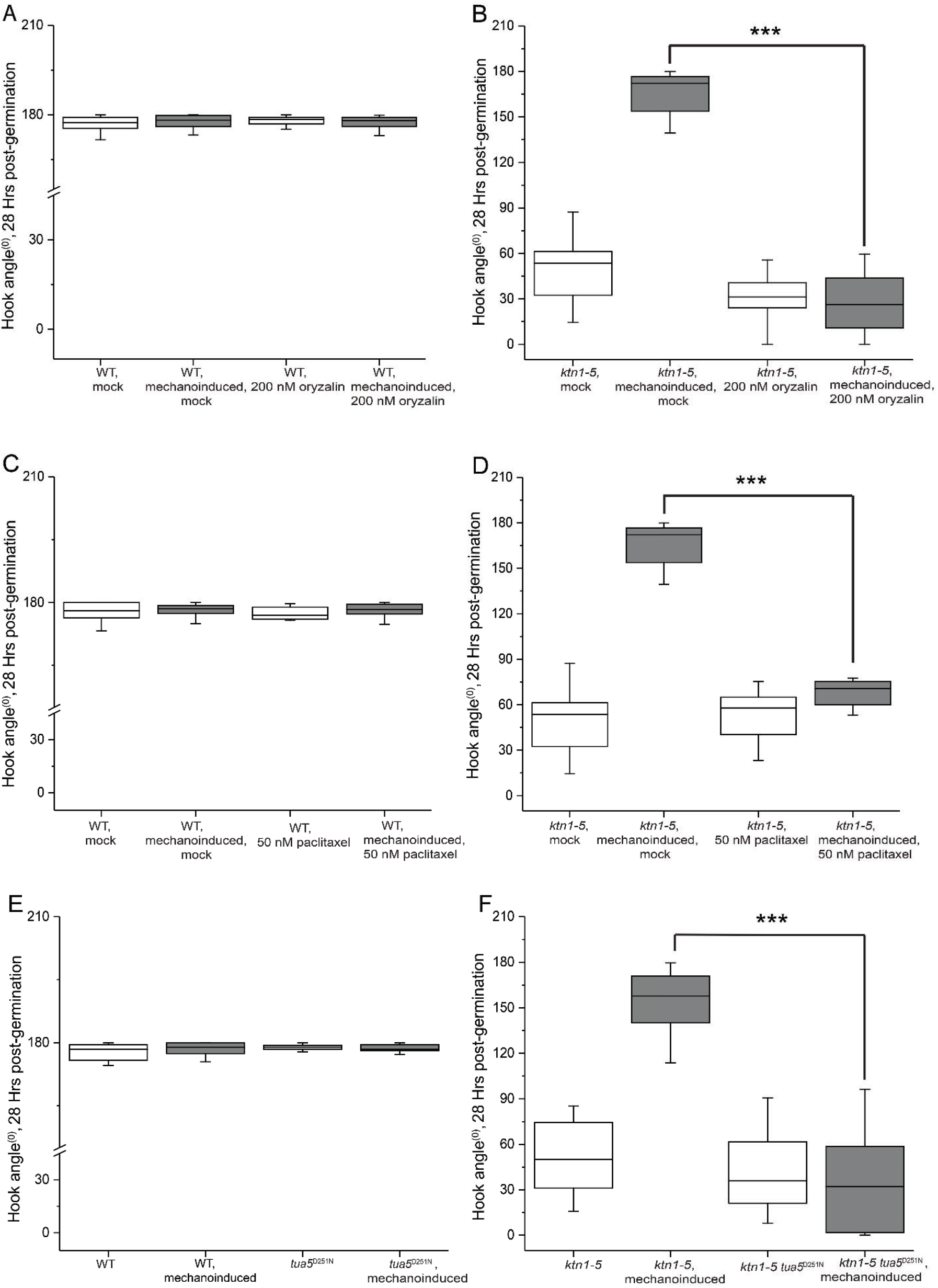
Chemical and genetic perturbations of microtubules eliminate mechanically-constrained hook formation in the *ktn1-5* mutant. Quantification of apical hook angles in WT and *ktn1-5* seedlings (in the presence and absence of mechanical constraint) treated with 200 nM oryzalin (A, B) or 50 nM paclitaxel (C, D) or in the presence of the *tua5*^D251N^ mutation (E, F), 28 Hrs post-germination. n = 18 seedlings per group, box plots showing the first and third quartiles, split by the median; whiskers extend to minimum and maximum values. Indicated groups are compared by Student’s two tailed t-test, asterisks (***) indicate P < 0.0001.

**Figure. S2.**
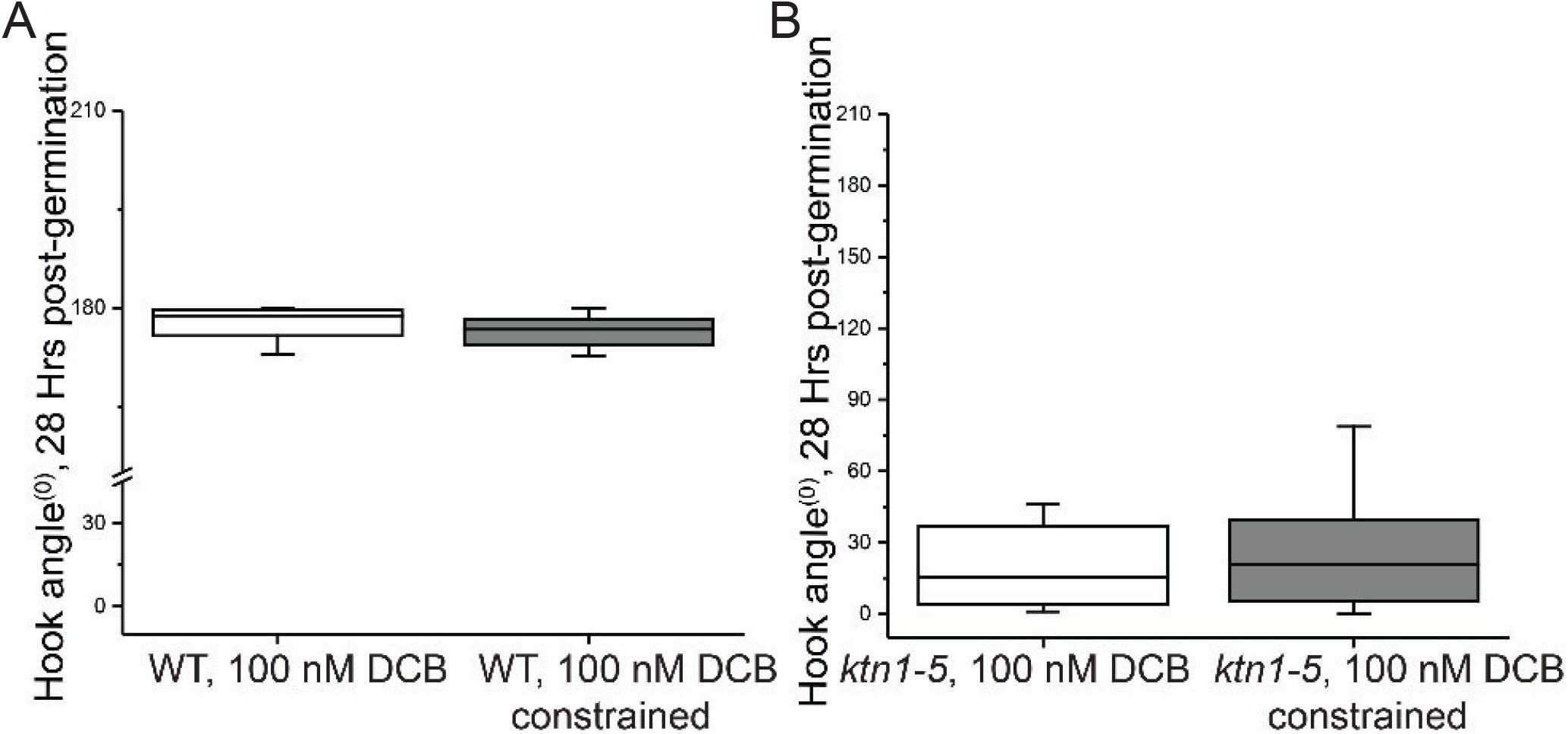
Mechanically constrained hook formation in *ktn1-5* is hypersensitive to the cellulose biosynthesis inhibitor DCB. Quantification of apical-hook angles in WT (A) and *ktn1-5* (B) treated with 100 nM DCB in the absence or presence of mechanical constraint, 28 Hrs after germination. n = 16 seedlings/ group.

**Figure. S3.**
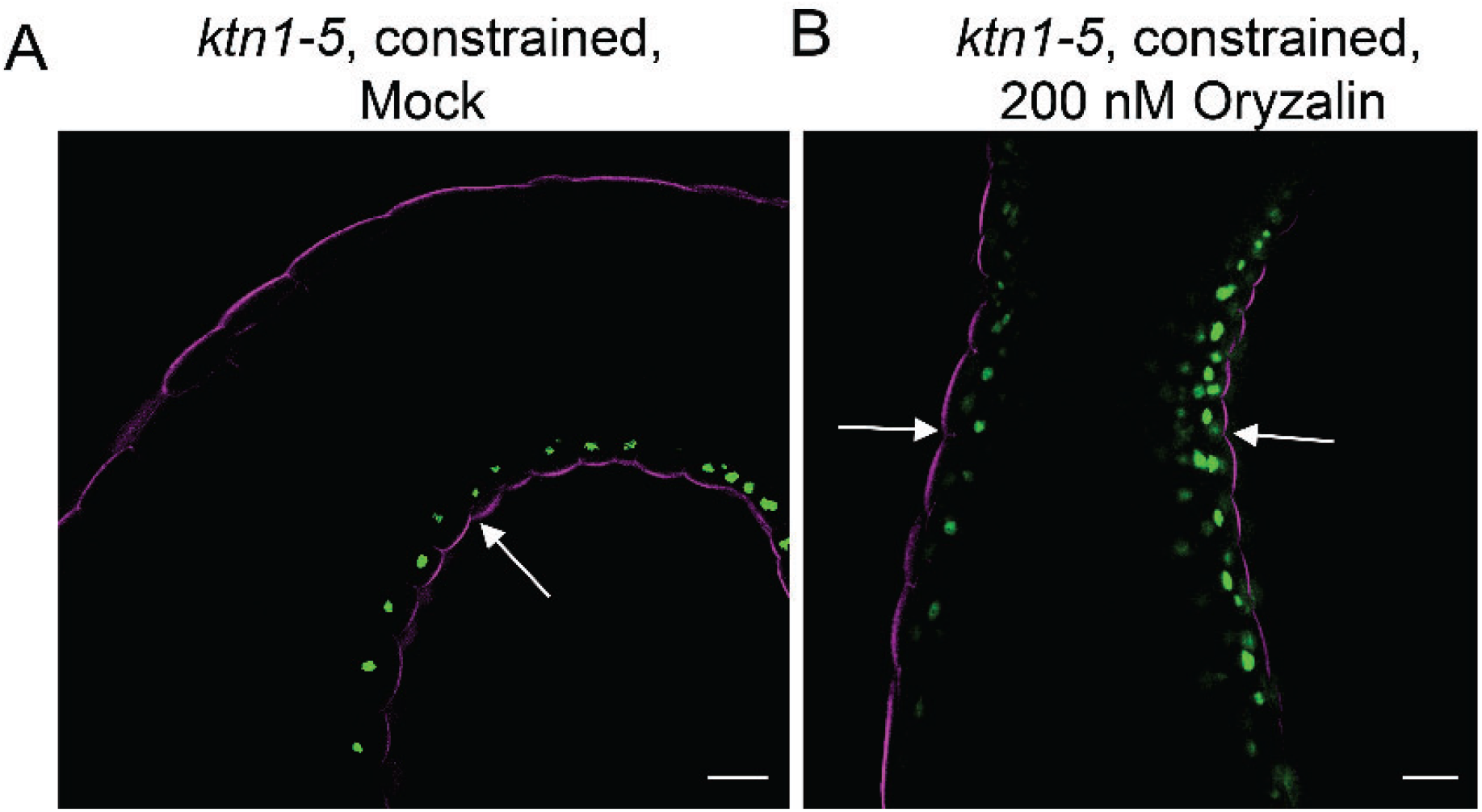
Representative image of DR5:: Venus-N7 signal in mechanically constrained *ktn1-5* seedlings grown inside mock MS agar (A) or MS agar supplemented with 200 nM Oryzalin (B). The seedlings are counterstained with propidium iodide. DR5:: Venus-N7 signal is indicated by arrows. At least 10 seedlings were observed per group.

**Figure. S4.**
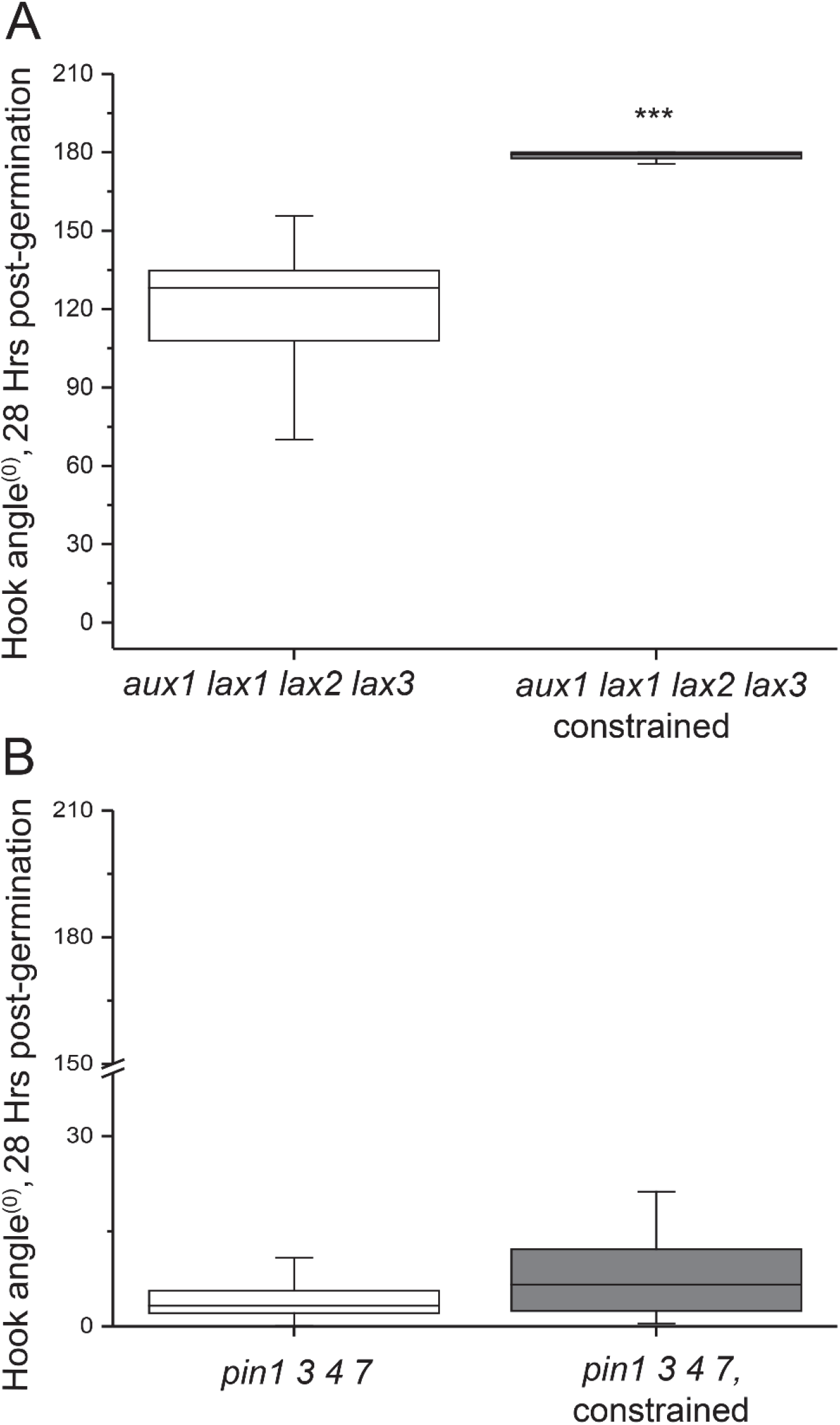
Involvement of auxin carriers in mechanically constrained hook formation. Quantification of apical-hook angles in *aux1 lax1 lax2 lax3* quadruple mutant (A) and *pin1 3 4 7* quadruple mutant (B) growing on the surface of MS agar or growing through the gel (constrained), 28 Hrs after germination, n = 16 seedlings/ group. Box plots showing the first and third quartiles, split by the median; whiskers extend to minimum and maximum values. Indicated groups are compared by Student’s two tailed t-test, asterisks (***) indicate P < 0.0001.

**Figure. S5.**
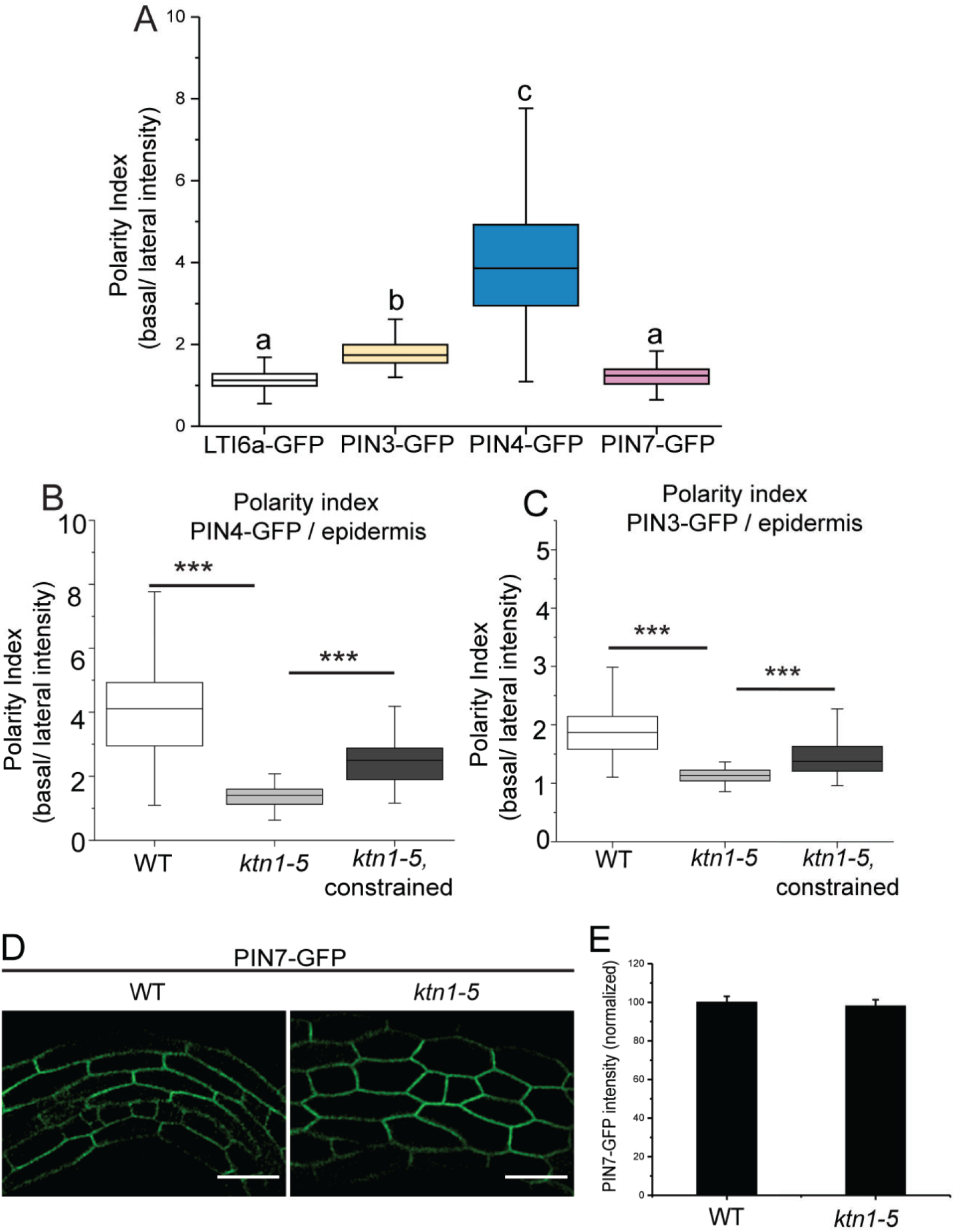
Polarity of auxin transporters. (A) Quantification of polarity index (basal/ lateral fluorescence intensity) of PIN3-GFP, PIN4-GFP, PIN7-GFP and LTI6a-GFP in hook epidermal cells. Polarity index was measured from at least 100 cells from 10 seedlings/ group. Polarity of PINs are compared with the nonpolar localization of LTI6a-GFP by one-way ANOVA with Tukey’s post-hoc test. Groups with significant difference (P < 0.05) are indicated by different letters. (B) Quantification of polarity index (basal/ lateral fluorescence intensity) of PIN4-GFP (A) and PIN3-GFP (B) in epidermal cells (related to Figure. 4A and 4B). Polarity index was measured from at least 100 cells from 10 seedlings/ group. Box plots show the first and third quartiles, split by the median; whiskers extended to minimum and maximum values. Asterisks (***) indicate P < 0.0001, ns = non-significant at P ≤ 0.05 (Student’s two tailed t-test). (C) Localization of PIN7-GFP in WT and ktn1-5 hook region. (D) quantification of PIN7-GFP intensities in WT and ktn1-5 hook. At least 100 cells from 10 seedlings per group was measured. Bars represent normalized mean ± S.E.M, ns = non-significant at P ≤ 0.05 (Student’s two tailed t-test).

**Figure. S6.**
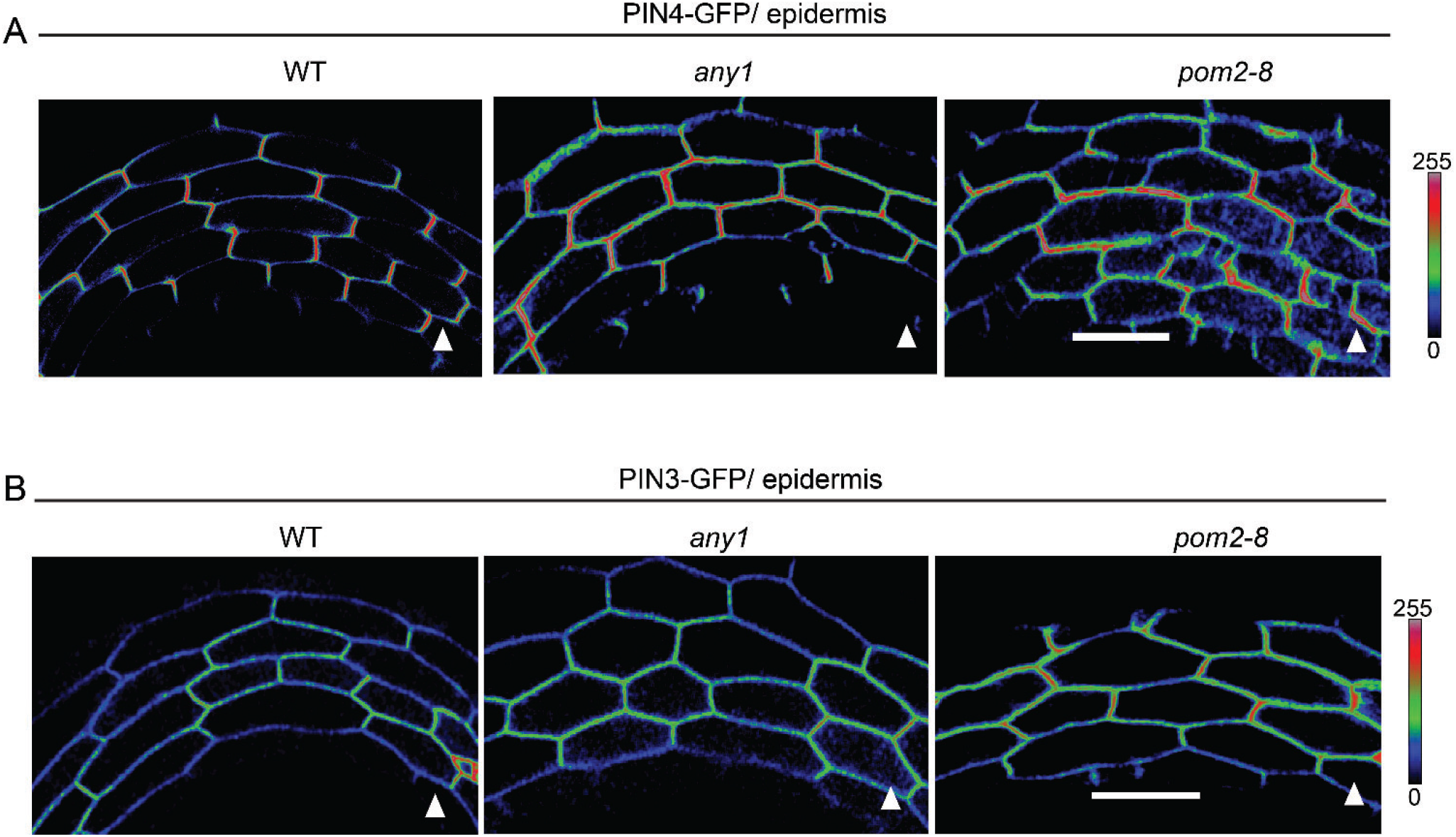
Representative images of PIN4-GFP (A), PIN3-GFP (B) in epidermal cells of WT, *any1* and *pom2-8* apical hook. Triangles denote the end of the hypocotyl with the shoot apical meristem. Scale bar = 50 μm. Color bars in (A) and (B) represent 8-bit intensity range of 0-255.

**Figure. S7.**
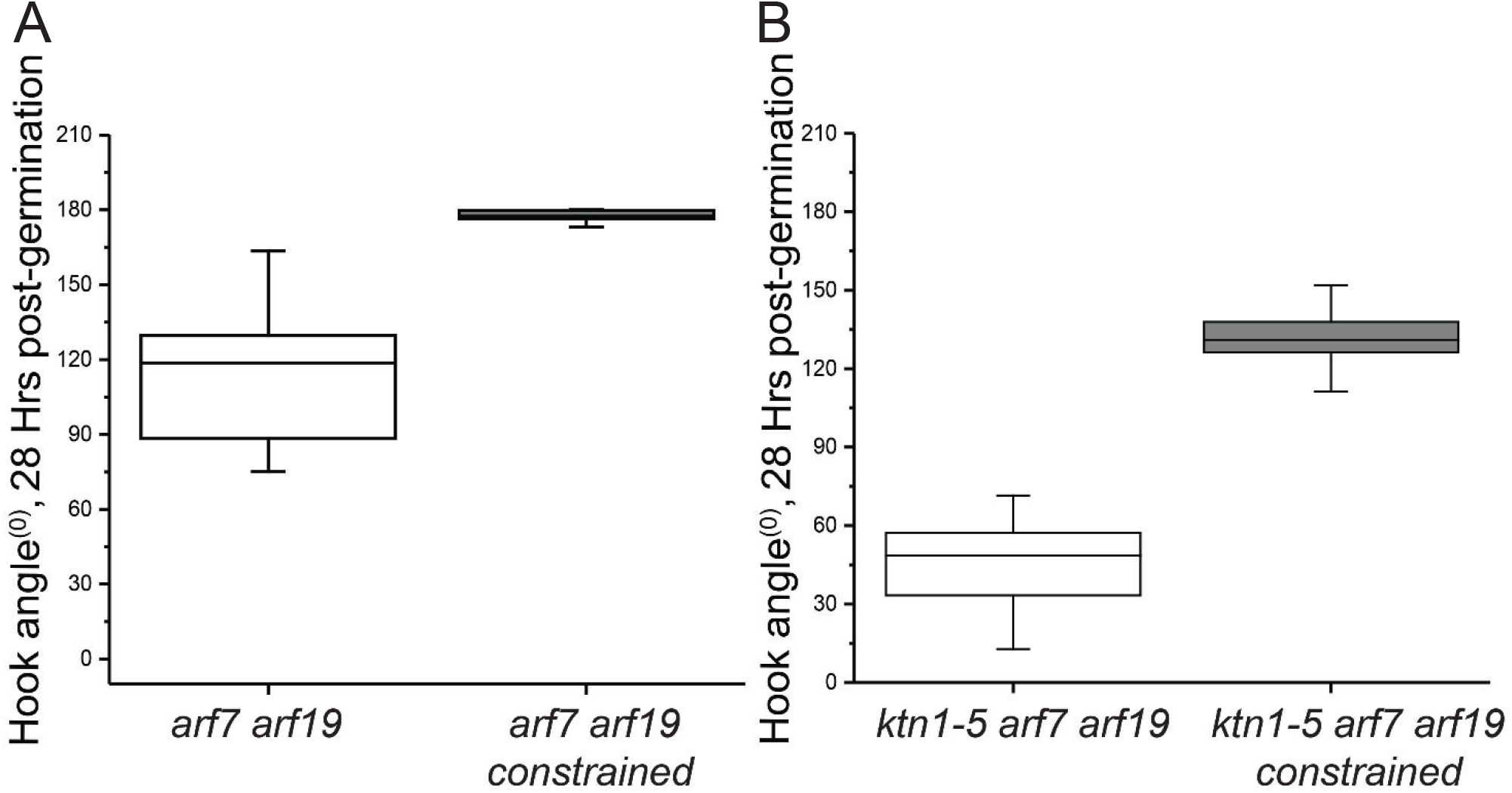
Mechanically constrained hook formation is not dependent on ARF7 and ARF19. Quantification of apical-hook angles in *arf7 arf19* (A) and *ktn1-5 arf7 arf19* (B) seedlings. growing on the surface of MS agar or growing through the gel (constrained), 28 Hrs after germination, n = 16 seedlings/ group. Box plots showing the first and third quartiles, split by the median; whiskers extend to minimum and maximum values.

## Notes

#### Summary of Updates

We have updated the the suppl data

